# Cell type-specific regulation by different cytokinetic pathways in the early embryo

**DOI:** 10.1101/2024.06.27.601054

**Authors:** Caroline Q. Connors, Sophia L. Martin, Julien Dumont, Mimi Shirasu-Hiza, Julie C. Canman

## Abstract

Cytokinesis, the physical division of one cell into two, is typically assumed to use the same molecular process across animal cells. However, regulation of cell division can vary significantly among different cell types, even within the same multicellular organism. Using six fast-acting temperature-sensitive (ts) cytokinesis-defective mutants, we found that each had unique cell type-specific profiles in the early *C. elegans* embryo. Certain cell types were more sensitive than others to actomyosin and spindle signaling disruptions, disrupting two members of the same complex could result in different phenotypes, and protection against actomyosin inhibition did not always protect against spindle signaling inhibition.

## Description

Cytokinesis, the physical division of one cell into two daughter cells, is critical for multicellular organismal development and tissue homeostasis. On one hand, defects in cytokinesis can lead to diseases such as cancer (Lacroix & Maddox, 2012; Lens & Medema, 2019). On the other hand, in some cell types (*e.g*., cardiomyocytes, hepatocytes, etc.), cytokinesis is naturally programmed to fail, leading to binucleation and polyploidy (Lacroix & Maddox, 2012). In animal cells, cytokinesis is driven by constriction of an actomyosin contractile ring at the cell division plane, positioned by signaling from spindle microtubules (for review see (D’Avino et al., 2015; Green et al., 2012)) and cell polarity (Cabernard et al., 2010; Jordan et al., 2016). The mechanisms that underlie cell type-specific regulation of cytokinesis are not well understood.

Here we compared the cell type-specific regulation of different cytokinesis pathways in the same model system. The early *C. elegans* lineage map is well characterized and invariant from embryo to embryo, making each cell easy to identify (**Fig. 1A**) (Sulston et al., 1983). Asymmetric cleavage divisions and cell fate signaling lead to differentiation and cell type-specific fate specification as early as the 2-cell stage. Thus, the early worm embryo serves as a simple multicellular system to study cell type-specific responses to genetic perturbations in cytokinesis in the same organism.

**Figure 1:**
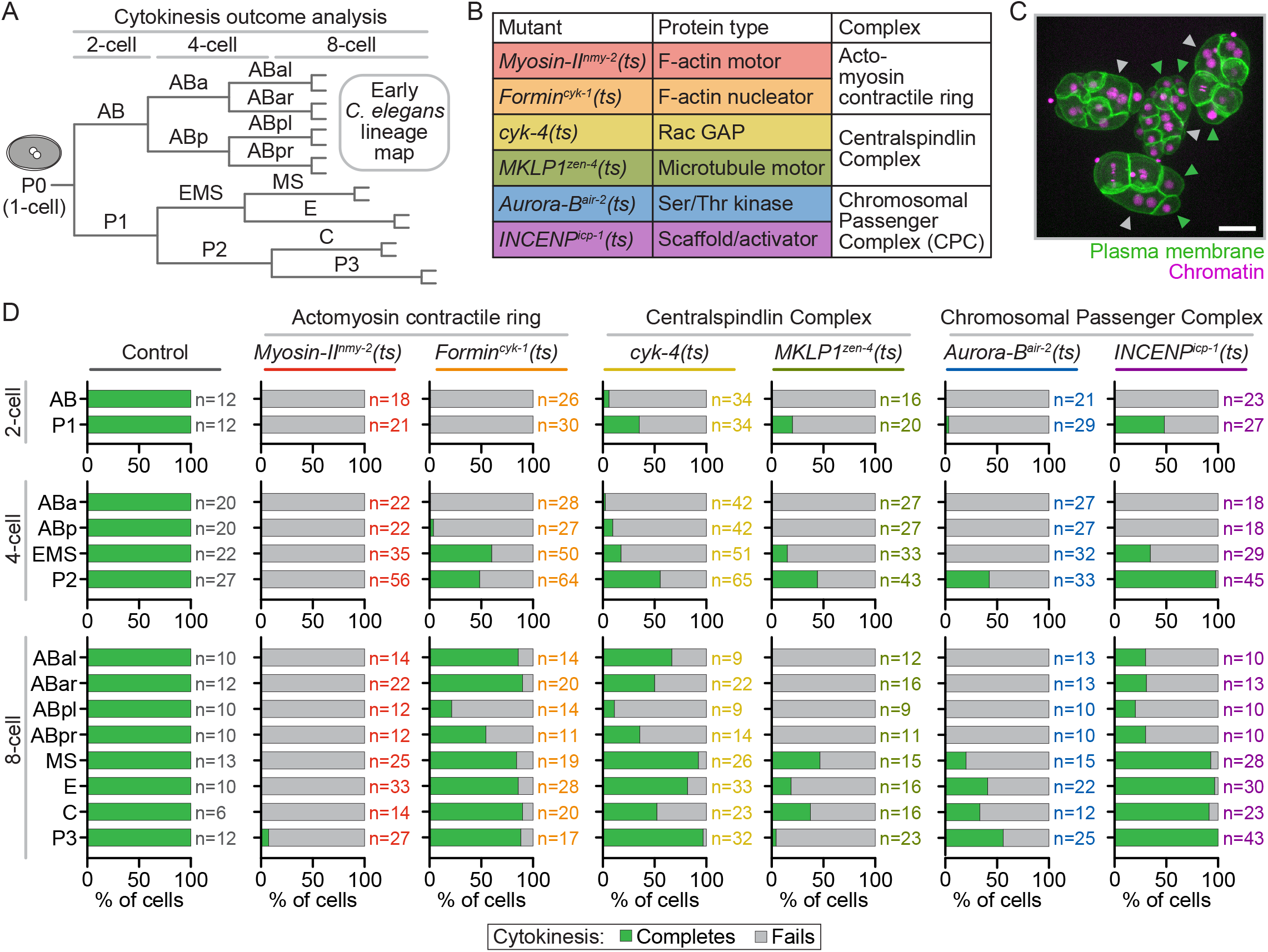
Cytokinesis outcomes in the early *C. elegans* embryo in different temperature sensitive mutants. **A)** Schematic depicting the early *C. elegans* embryonic lineages from the 1-cell to 8-cell stage. **B)** Temperature-sensitive (ts) mutant alleles, protein functions, and complexes affected. **C)** Representative maximum projection image of clustered embryos scored for cytokinesis completion or failure. Scale bar=20 μm; green arrowheads=cytokinesis completes; grey arrowheads=cytokinesis fails. **D)** Graphs showing the percentage of each cell type that completes (green) or fails (gray) in cytokinesis after upshift to restrictive temperature by column in control (no ts mutations), *myosin-II*^*nmy-2*^*(ts)* and *formin*^*cyk-1*^*(ts)* (actomyosin contractile ring), *cyk-4(ts)* and *MKLP1*^*zen-4*^*(ts) (*Centralspindlin complex), and *Aurora-B*^*air-2*^*(ts)* and *INCENP*^*icp-1*^*(ts)* (Chromosomal Passenger Complex (CPC)) mutant 2-cell (top rows), 4-cell (middle rows), and 8-cell (bottom rows) embryos. n=number individual cells scored per cell type and genotype (by color) indicated to the right of each bar.

To specifically disrupt cytokinesis protein function in individual cells during early development, we turned to our collection of fast-acting (≤20 sec) temperature-sensitive (ts) cytokinesis-defective mutants. We used six ts mutants that affect three different cytokinesis complexes (Davies et al., 2014): the actomyosin contractile ring (*myosin-II*^*nmy-2*^*(ts)* and diaphanous *formin*^*cyk-1*^*(ts)*); centralspindlin complex (*MKLP1*^*zen-4*^*(ts)* and *cyk-4(ts)*); and the Chromosomal Passenger Complex (CPC) (*INCENP*^*icp-1*^*(ts)* and *Aurora-B*^*air-2*^*(ts)*) (**Fig. 1B**) (Canman et al., 2008; Davies et al., 2014; Liu et al., 2010; Severson et al., 2000). These conditional mutations allow cell division to occur at permissive temperature, but completely block cytokinesis in the 1-cell embryo when upshifted to restrictive temperature (Davies et al., 2014).

To test for cell type-specific responses to acute cytokinesis complex disruption in *C. elegans*, we imaged control and ts mutant embryos before and after upshifting to restrictive temperature (**Fig. 1C**). To control temperature, we used the Therminator, a fluidic device that allows rapid (<17 sec) upshift from permissive (16ºC) to restrictive temperature (26ºC) while simultaneously imaging using a spinning disc confocal microscope (Davies et al., 2014; Davies et al., 2017). We examined cytokinesis success or failure in each cell in 2-cell through 8-cell stage embryos after upshift to restrictive temperature before either anaphase onset (actomyosin and centralspindlin mutants, controls) or nuclear envelope breakdown (CPC mutants) (Davies et al., 2014) (**Fig. 1D**). As expected, in control 2-thru 8-cell embryos, all cells completed cytokinesis successfully upon temperature upshift and, in *myosin-II*^*nmy-2*^*(ts)* embryos, almost all cells failed in cytokinesis (**Fig. 1D**). In all other ts mutants, we observed cell type differences starting at the 2-cell or 4-cell stage (**Fig. 1D**). In *formin*^*cyk-1*^*(ts)* 2-cell embryos, both cells failed in cytokinesis, but in 4-cell embryos, dramatic cell type-specific differences in the outcome of cytokinesis was observed: ABa and ABp failed in cytokinesis but EMS and P2 frequently completed cytokinesis, as described (Connors et al., 2024; Davies et al., 2018). In *formin*^*cyk-1*^*(ts)* 8-cell embryos, most cells completed cytokinesis >80% of the time (except for ABpl and ABpr; 21% and 55% completed cytokinesis, respectively). In centralspindlin and CPC mutant 2-cell embryos and 4-cell embryos, a similar pattern of cytokinesis failure or completion were observed in each cell type (**Fig. 1D**): in 2-cell embryos, the P1 cell completed cytokinesis at a higher rate than the AB cell (except in CPC *Aurora-B*^*air-2*^*(ts)* embryos), and in 4-cell embryos, the P2 cell frequently (42-98%) completed cytokinesis whereas EMS only completed cytokinesis 0-34% of the time. In 8-cell embryos, there were cell type-specific differences in cytokinesis for all centralspindlin and CPC mutants, even for mutations that affect the same protein complex (**Fig. 1D**). For example, in centralspindlin *cyk-4(ts)* embryos at the 8-cell stage, most cells completed cytokinesis, except for ABpl and ABpr; in contrast, in centralspindlin *MKLP1*^*zen-4*^*(ts)* 8-cell embryos, only MS, E, and C were able to complete cytokinesis at any frequency (**Fig. 1D**). Additionally, in CPC *Aurora-B*^*air-2*^*(ts)* 8-cell embryos, no anterior cell types (ABal, ABar, ABpl, ABpr) were able to complete cytokinesis while all posterior cell types (MS, E, C, P3) completed cytokinesis 20-56% of the time (**Fig. 1D**). In contrast, in CPC *INCENP*^*icp-1*^*(ts)* 8-cell embryos, cells in the anterior (ABal, ABar, ABpl, ABpr) were able to complete cytokinesis 20-31% of the time, while cells in the posterior (MS, E, C, P3) completed cytokinesis a striking 91-100% of the time, (**Fig. 1D**). Thus, cell type-specific regulation of cytokinesis differs after disruption of different protein complexes/activities and can be widely observed in the early worm embryo.

We were initially surprised that the pattern of cell type-specific cytokinesis success versus failure was so different between cytokinesis-defect mutants that affect the same complex. Yet each mutant differentially affects protein function and complex activity. The formin^CYK-1^ ts mutation affects the dimerization domain and greatly compromises F-actin nucleation activity (Davies et al., 2014); our previous results suggest that cell fate determinants regulate cell type-specific responses to formin^CYK-1^ inactivation (Connors et al., 2024; Davies et al., 2018). The myosin-II^NMY-2^ mutation is in the neck region (Liu et al., 2010), which likely decouples and inactivates myosin motor domains. Importantly, myosin-II^NMY-2^ plays an essential role in cell polarity/asymmetric cell division and cell fate specification/cell identity (Guo & Kemphues, 1996; Liu et al., 2010); thus, this mutant will reduce the differences between cell types and affect cytokinesis in all cells equally, as was observed (**Fig. 1D**). The centralspindlin complex mutants also have dramatically different effects on complex activity. While the MKLP1^ZEN-4^ ts mutation completely blocks centralspindlin complex formation and central spindle microtubule bundling (Pavicic-Kaltenbrunner et al., 2007; Severson et al., 2000), the CYK-4 ts mutation only affects Rac GAP (GTPase-activating protein) activity (Canman et al., 2008). Similarly, while the Aurora-B^AIR-2^ ts mutation disrupts the kinase domain and thus catalytic activity of the CPC (Bishop & Schumacher, 2002), the INCENP^ICP-1^ ts mutation reduces but does not totally block Aurora-B^AIR-2^ kinase activity (Davies et al., 2014). Thus, it seems logical that cytokinesis is more frequently successful in the less disruptive mutant condition for each complex.

We also observed cases of “inherited” resistance to disruption of cytokinesis proteins from mother cells to daughter cells, as well as *de novo* resistance in daughter cells born from mother cells that typically would have failed in cytokinesis upon protein inactivation. In general, most cells in the 8-cell embryo completed cytokinesis successfully after cytokinesis protein inactivation more often than their mother cells in the 4-cell embryo, independent of genotype. As an extreme example, in *cyk-4(ts)* 4-cell embryos, while the ABa cell almost always failed in cytokinesis, both of its 8-cell stage daughter cells, ABal and ABar, completed cytokinesis at a high frequency (67% and 50% cytokinesis completion in ABal and ABar, respectively) (**Fig. 1D**). Still, we noted specific cells in specific genotypes that divided *less* frequently than their mother cells. For example, in *MKLP1*^*zen-4*^*(ts)* 4-cell embryos, the P2 cell successfully divided 44% of the time but this resistance was only inherited by one daughter cell and not the other (38% and 4% cytokinesis completion in C and P3 daughter cells, respectively) (**Fig. 1D**). Future experiments will determine if similar or different mechanisms drive cell type-specific regulation of cytokinesis in different lineages and how cell fate specification regulates the cytokinesis machinery to accommodate the development and function of that cell within the context of a multicellular organism.

## Methods

### Worm husbandry

Worm strains were grown on 60 mm petri plates (T3308, Tritech) filled with (PourBoy 4, Tritech) 10.5 mL nematode growth media (NGM) (23 g Nematode Growth Medium (Legacy Biologicals), 1 mL 1M CaCl_2_, 1 mL 1M MgSO_4_, 25 mL 1M K_3_PO_4_, 975 mL ddH_2_O) seeded with 500 μL *E. coli* (OP50), as in (Brenner, 1974). Strains were maintained at 16ºC in incubators (Binder). We note that wormbase.org was used as a resource throughout this work (Sternberg et al., 2024).

### Embryo preparation for imaging

On imaging days, gravid young adult hermaphrodites were maintained in an incubator at 13-14ºC (2720213W, Wine Enthusiast) and dissected on a stereo microscope (Olympus SZX16 with SDF PLAPO 1XPF objective) in cooled (13-14ºC) M9 buffer in a watch glass (742300, Carolina Biologicals). A hand-pulled glass pipette (VWR Pasteur Pipette) or a borosilicate glass capillary (World Precision Instruments) was used as a mouth pipette to transfer embryos onto a thin 2% agar pad placed on top of the Therminator specimen holder (Davies et al., 2014; Davies et al., 2017). Embryos were clustered using a single hair tool (Ted Pella). A 30 mm round No. 1.5 glass coverslip (Bioptechs) was mounted on top of the embryos for imaging.

### Time-lapse live cell imaging

Time lapse live cell imaging was done on an inverted microscope stand (Nikon, Eclipse Ti) with a spinning disc confocal head (CSU-10, Yokogawa; Borealis upgrade (Spectral Applied Research)), a CCD camera with 2 x 2 binning (Orca-R2, Hamamatsu), a Piezo-motorized stage (ASI) for Z-sectioning, and either a 20x Plan Apo 0.75 N.A. dry objective (Nikon) or a 40x Plan Apo Lambda 0.95 N.A. dry objective (Nikon). An acousto-optic tunable filter (Spectral Applied Research) was used to control excitation laser light (150 mW 488 nm (GFP) and 561 nm (mCherry); ILE-2, Spectral Applied Research) and a filter wheel (Sutter) was used for 525/50 nm or 620/60 nm emission filter (Chroma) selection. Focus was maintained (Nikon, Perfect Focus) before each timepoint: 12 x 2 or 11 x 2 μm Z-sections every 60 or 90 seconds; 100 ms and 100-150 ms exposures for GFP and mCherry channels, respectively.

### Temperature control for live cell imaging

Live cell imaging was performed in an imaging room with a mini-split heat pump to control temperature (MHWX, MultiAqua). Room temperature (19-23ºC) was continuously monitored with 4-5 digital thermometers and a Bluetooth sensor (SensorPush) on the microscope stage. The Therminator was used as described previously (Davies et al., 2014; Davies et al., 2017). Briefly, one water/isopropanol bath was set to permissive temperature (16ºC) and the second bath was set to restrictive temperature (26ºC). 2-cell through 8-cell embryos were maintained on the specimen holder at 16ºC until the desired time at which point the Therminator was switched to use the 26ºC water bath and rapidly upshift sample temperature to restrictive temperature (25.5-27.3ºC) prior to anaphase onset (control, *myosin-II*^*nmy-2*^*(ts), formin*^*cyk-1*^*(ts), MKLP1*^*zen-4*^*(ts), and cyk-4(ts)* embryos) or nuclear envelope breakdown (*INCENP*^*icp-1*^*(ts)* and *Aurora-B*^*air-2*^*(ts)* embryos) in each cell.

### Cytokinesis outcome analysis

FIJI (FIJI is Just ImageJ) software (Schindelin et al., 2012) was used for all data analysis. Cytokinesis outcomes were scored manually on maximum projection images of both channels (GFP::PH^pLCδ^ and mCherry::histone H2B^HIS-58^) as in (Connors et al., 2024). Individual cells were scored only if upshift to restrictive temperature occurred before anaphase onset (or nuclear envelope breakdown (≥450 seconds prior to anaphase onset) in *INCENP*^*icp-1*^*(ts)* and *Aurora-B*^*air-2*^*(ts)* embryos) in that cell and at least one of its daughter cells entered anaphase before the end of the image series.

## Author Contributions

CQC and JCC conceived of and designed experiments. CQC and SLM performed the experiments. CQC and JCC made the figure. CQC, SLM, JD, MSH, and JCC made significant intellectual contributions and helped write (or edit) the manuscript.

## Acknowledgements

We thank all members of the Canman, Dumont, and Shirasu-Hiza labs for helpful discussions and support. We thank Sophia Tony-Egbuniwe, Eva Sophia Blake, Adriana Hernandez, and Michelle (Mimi) Schmidt for making worm plates and other critical lab reagents. We are grateful to Karen Oegema, Craig Mello, Bruce Bowerman, and the CGC (funded by NIH Office of Research Infrastructure Programs (P40 OD010440)) for providing worm strains. This work was funded by: NIH R01GM117407 (JCC), R01GM130764 (JCC), European Research Council CoG ChromoSOMe N°819179 (JD), NIH R01AG045842 (MSH), and NIH R35GM127049 (MSH). The authors declare no competing financial interests.

## Abbreviations

ts: temperature sensitive
F-actin: filamentous actin
GAP: GTPase activating protein
MKLP-1: mitotic kinesin-like protein-1
ZEN-4: zygotic enclosure-defective
CYK-1/-4: cytokinesis-defective-1/-4
CPC: chromosomal passenger complex
INCENP: inner centromere protein
ICP-1: INCENP homolog
NMY-2: non-muscle myosin-II

