## Supplementary material for "Cell type-specific regulation by different cytokinetic pathways in the early embryo": Table S1

**Connors et al Table S1: Worm strains table**

| Strain Name | Description | Full Genotype | Reference (first use) |
|---|---|---|---|
| OD95 | control | *unc-119(ed3)* ltIs38 [pAA1; pie-1/GFP::PH(PLC1delta1); unc-119 (+)] III; ltIs37 [pAA64; pie-1/mCHERRY::his-58; unc-119 (+)] IV* | Audhya *et al.* , 2005, *J. Cell Biology* |
| JCC637 | *nmy-2(ne3409ts)* mutant | *nmy-2(ne3409ts)I; unc-119(ed3)* ltIs38 [pAA1; pie-1/GFP::PH(PLC1delta1)III; unc-119 (+)]; ltIs37 [pAA64; pie-1/mCHERRY::his-58; unc-119 (+)]IV* | Liu *et al.* , 2010, *Dev. Biol.* |
| JCC146 | *cyk-1(or596ts)* mutant | *cyk-1(or596ts) unc-119(ed3)* ltIs38 [pAA1; pie-1/GFP::PH(PLC1delta1); unc-119 (+)] III; ltIs37 [pAA64; pie-1/mCHERRY::his-58; unc-119 (+)] IV* | Davies *et al.* , 2014, *Dev. Cell* |
| OD239 | *cyk-4(or749ts)* mutant | *cyk-4(or749ts) unc-119(ed3)* ltIs38 [pAA1; pie-1/GFP::PH(PLC1delta1)III; unc-119 (+)]; ltIs37 [pAA64;pie-1/mCHERRY::his-58; unc-119 (+)]IV* | Canman *et al.* , 2008, *Science* |
| JCC754 | *zen-4(or153ts)* mutant | *unc-119(ed3)* ltIs38 [pAA1; pie-1/GFP::PH(PLC1delta1)III; unc-119 (+)]; zen-4(or153ts) ltIs37 [pAA64;pie-1/mCHERRY::his-58; unc-119 (+)]IV* | Severson *et al.* , 2000, *Curr. Biol.* ; Pavicic-Kaltenbrunner *et al.* , 2007, *Mol. Biol. Cell* |
| JCC179 | *air-2(or207ts)* mutant | *air-2(or207ts)I; unc-119(ed3)* ltIs38 [pAA1; pie-1/GFP::PH(PLC1delta1)III; unc-119 (+)]; ltIs37 [pAA64;pie-1/mCHERRY::his-58; unc-119 (+)]IV* | Severson *et al.* , 2000, *Curr. Biol.* |
| JCC030 | *icp-1(or663ts)* mutant | *icp-1(or663ts)I; unc-119(ed3)* ltIs38 [pAA1; pie-1/GFP::PH(PLC1delta1)III; unc-119 (+)]; ltIs37 [pAA64;pie-1/mCHERRY::his-58; unc-119 (+)]IV* | Davies *et al.* , 2014, *Dev. Cell* |

*NOTE: The *unc-119(ed3* ) mutation may be in this genetic background but not visible due to rescuing *unc-119(+)* sequence(s). The endogenous *unc-119* locus has not been sequenced.
